# Discovery of the role of a SLOG superfamily biological conflict systems associated protein IodA (YpsA) in oxidative stress protection and cell division inhibition in Gram-positive bacteria

**DOI:** 10.1101/451617

**Authors:** Robert S. Brzozowski, Gianni Graham, A. Maxwell Burroughs, Mirella Huber, Merryck Walker, Sameeksha S. Alva, L. Aravind, Prahathees J. Eswara

## Abstract

Bacteria adapt to different environments by regulating cell division and several conditions that modulate cell division have been documented. Understanding how bacteria transduce environmental signals to control cell division is critical to comprehend the global network of cell division regulation. In this article we describe a role for *Bacillus subtilis* YpsA, an uncharacterized protein of the SLOG superfamily of nucleotide and ligand-binding proteins, in cell division. We observed that YpsA provides protection against oxidative stress as cells lacking *ypsA* show increased susceptibility to hydrogen peroxide treatment. We found that increased expression of *ypsA* leads to cell division inhibition due to defective assembly of FtsZ, the tubulin-like essential protein that marks the sites of cell division. We showed that cell division inhibition by YpsA is linked to glucose availability. We generated YpsA mutants that are no longer able to inhibit cell division. Finally, we show that the role of YpsA is possibly conserved in Firmicutes, as overproduction of YpsA in *Staphylococcus aureus* also impairs cell division. Therefore, we propose *ypsA* to be renamed as *iodA* for <u>*i*</u>nhibitor <u>*o*</u>f <u>*d*</u>ivision.

**IMPORTANCE:** Although key players of cell division in bacteria have been largely characterized, the factors that regulate these division proteins are still being discovered and evidence for the presence of yet-to-be discovered factors has been accumulating. How bacteria sense the availability of nutrients and how that information is used to regulate cell division positively or negatively is less well-understood even though some examples exist in the literature. We discovered that a protein of hitherto unknown function belonging to the SLOG superfamily of nucleotide/ligand-binding proteins, YpsA, influences cell division in *Bacillus subtilis* by integrating metabolic status such as the availability of glucose. We showed that YpsA is important for oxidative stress response in *B. subtilis*. Furthermore, we provide evidence that cell division inhibition function of YpsA is also conserved in another Firmicute *Staphylococcus aureus*. This first report on the role of YpsA (IodA) brings us a step closer in understanding the complete tool set that bacteria have at their disposal to regulate cell division precisely to adapt to varying environmental conditions.

## INTRODUCTION

Cell division in bacteria is a well-orchestrated event that is achieved by the concerted action of approximately a dozen different key division proteins (1). Amongst them a protein central to cell division in most bacteria is the tubulin homolog, FtsZ, which marks the site of cytokinesis (2, 3). In addition to standard spatial regulators of septum positioning (4), factors that sense nutrient availability (5, 6), DNA damage (7-9), alternate external environment (10, 11), have been shown to influence cell division. The observation that cell division in model organisms *Escherichia coli* and *Bacillus subtilis* lacking well-studied Min and nucleoid occlusion regulatory systems undergo cell division largely unperturbed (12), prompted us to investigate the presence of other factors involved in cell division regulation. Here we describe the role of IodA (YpsA), a protein conserved in several members of the Firmicutes phylum.

The genes *iodA* (*ypsA*) and *gpsB* (formerly *ypsB*) are in a syntenous relationship in many Firmicute genomes (Fig. 1A). GpsB is a cell division protein that regulates peptidoglycan synthesis in *B. subtilis* (13, 14), *Streptococcus pneumoniae* (15, 16), and *Listeria monocytogenes* (17). More recently our group showed that *Staphylococcus aureus* GpsB affects the polymerization kinetics of FtsZ directly (18). As genes in a syntenous arrangement across multiple genomes, often referred to as conserved gene neighborhoods, are commonly indicative of functional relationships (19, 20), we were curious to study the function of YpsA in *B. subtilis*. Prior to our investigation, the crystal structure of *B. subtilis* YpsA was solved by a structural genomics group (PDB ID: 2NX2). Based on unique structure and sequence features (Fig. 1B), YpsA was classified as the founding member of the “YpsA proper” clade in the <u>*S*</u>MF/DprA/<u>*LOG*</u> (SLOG) protein superfamily (21). The SLOG superfamily contains a specific form of the Rossmannoid fold, and is involved in a range of nucleotide-related functions. These include the binding of low-molecule weight biomolecules, nucleic acids, free nucleotides, and the catalyzing of nucleotide-processing reactions (22-24). Recently, several members of the SLOG superfamily were further identified as key components in a newly-defined class of biological conflict systems centered on the production of nucleotide signals. In these systems, SLOG proteins are predicted to function either as sensors binding nucleotide signals or as nucleotide-processing enzymes generating nucleotide derivatives which function as signals (21). Despite these new reports, the precise function of YpsA and its namesake family have yet to be experimentally investigated.

**Figure 1.**
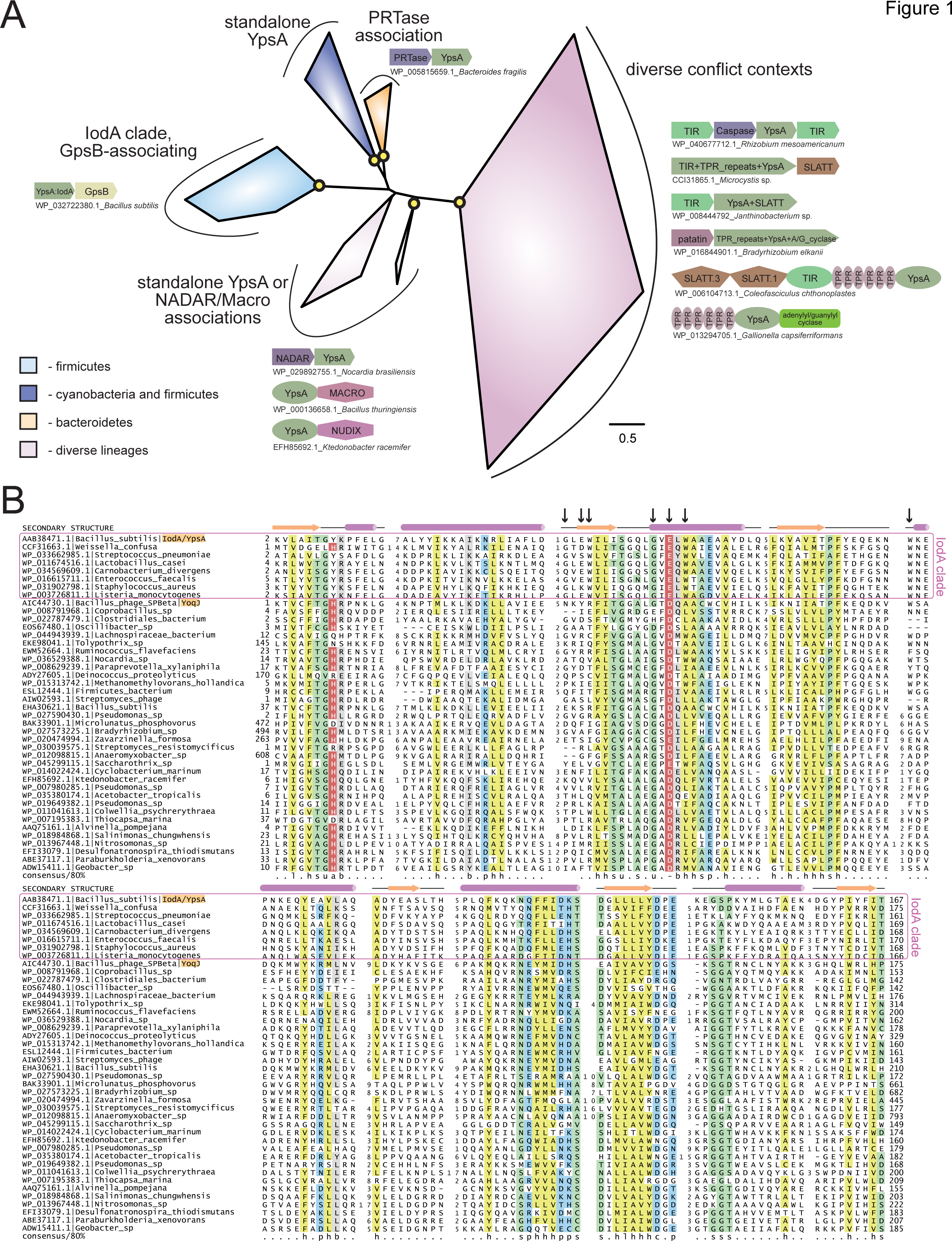
(A) Phylogenetic tree of the YpsA family, key branches with greater than 70% bootstrap support are denoted with yellow circles. Reproducible clades within the family are color-coded according to their phyletic distribution and labeled with names and representative conserved domain architectures and gene neighborhoods. For these genome context depictions, colored polygons represent discrete protein domains within a protein, while boxed arrows represent individual genes within a neighborhood. Each context is labeled with NCBI accession and organism name, separated by an underscore. For gene neighborhoods, the labeled gene contains the YpsA domain. Abbreviations: A/G_cyclase, adenylyl/guanylyl cyclase. (B) Multiple sequence alignment of the YpsA family of proteins. Secondary structure and amino acid biochemical property consensus are provided on the top and bottom lines, respectively. Black arrows at top of alignment denote positions subject to site-directed mutagenesis. Sequences are labeled to left with NCBI accession and organism name separated by vertical bars. Gene names from the text are provided after organism name, shaded in orange. Selected members of the IodA (YpsA) clade, which associate with GpsB, are enclosed in a purple box. Alignment coloring and consensus abbreviations as follows: b, big and gray; c, charged and blue; h, hydrophobic and yellow; l, aliphatic and yellow; p, polar and blue; s, small and green; u, tiny and green. The conserved aromatic position in the first loop, abbreviated ‘a’, and the conserved negatively‐charged position in the second helix, abbreviated ‘‐‘, are both colored in red with white lettering to distinguish predicted, conserved positions located within the active site pocket.

Here we report that (i) YpsA provides protection against oxidative stress; (ii) overexpression of *ypsA* causes mislocalization of FtsZ-GFP that results in cell filamentation which is dependent on glucose availability; (iii) YpsA-GFP forms dynamic foci that is likely mediated by nucleotide binding; and finally (iv) overexpression of *ypsA* in *S. aureus* results in cell enlargement, typical of cell division inhibition in cocci (25), suggesting a conserved function of YpsA across Firmicutes with very different cell-morphologies. In sum, these results constitute the first report on YpsA and its role in oxidative stress response and cell division regulation. Therefore, we propose to rename YpsA as IodA (<u>**i**</u>nhibitor <u>**o**</u>f <u>**d**</u>ivision) to best describe the function of YpsA.

## RESULTS

### YpsA provides oxidative stress protection

As a first step to study the significance of YpsA, we studied the phenotype of *ypsA* null strain in several stress-inducing conditions through standard disc-diffusion assay. As shown in Fig. 2A, we noticed that *ypsA* null cells exhibited a larger zone of inhibition in comparison to WT when incubated with discs soaked in 1 M H_2_O _2_(WT: 1.8 ± 0.45 mm; Δ*ypsA*: 7.6 ± 0.54 mm). It is noteworthy that *ypsA* transcript level is elevated upon hydrogen peroxide treatment (26, 27). To further evaluate this phenotype, we monitored the cells grown in liquid culture in the absence or presence of 1 mM H_2_O_2_ using fluorescence microscopy (Fig. 2B). Untreated cells lacking *ypsA* appear morphologically similar to WT. Although WT cells were tolerant to H_2_O_2_ treatment, Δ *ypsA* cells displayed obvious signs of “sick cells” en route to lysis such as membrane thickening, cell morphology change, condensed DNA (Fig. 2B; compare right top and middle panels). Quantification of H_2_O_2_-treated cells revealed that 27% of WT and 79% of Δ*ypsA* cells were sick (n =100). To test if this phenotype is specifically due to absence of YpsA, we introduced an inducible copy of *ypsA* at an ectopic locus. In the presence of inducer, H_2_O_2_-treated cells resemble WT (Fig. 2B, bottom panel; 23% sick cells, n =100) indicating that YpsA is responsible for providing protection against H_2_O_2_-induced oxidative stress.

**Figure 2.**
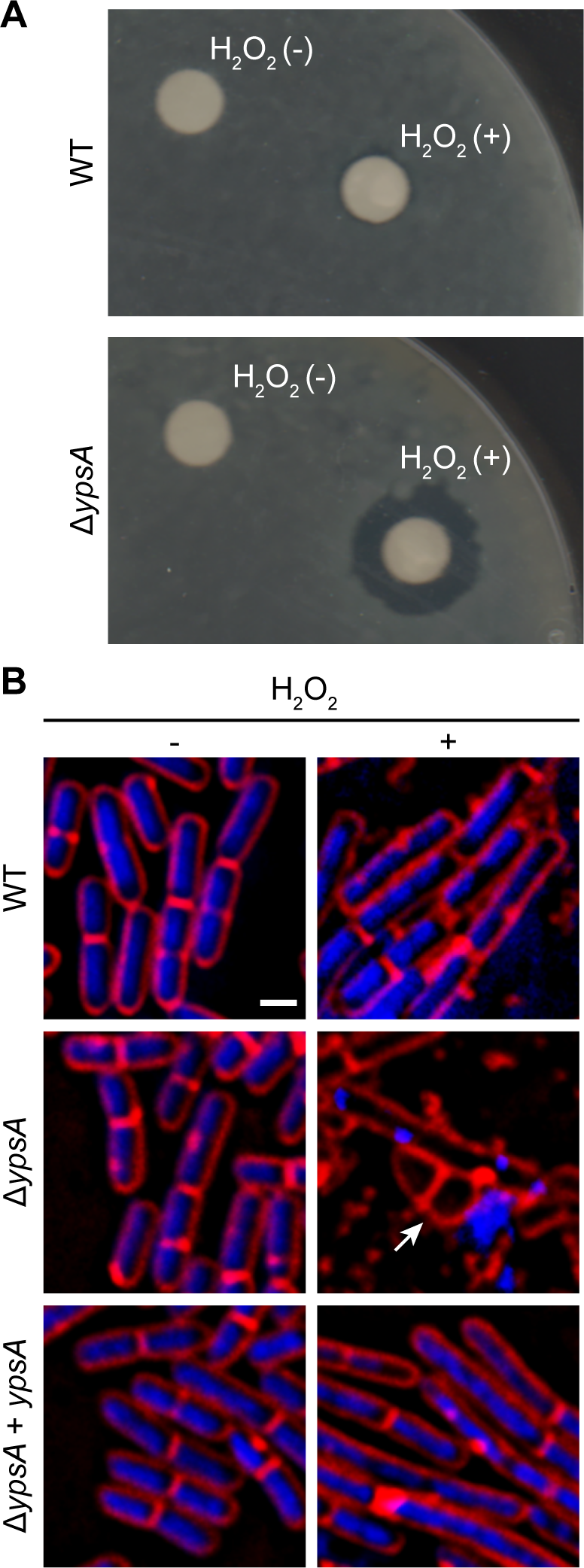
YpsA plays a role in oxidative stress response. (A) Disc diffusion assay with lawns made of WT (PY79) or a strain lacking *ypsA* (RB42) treated with blank disc and 1 M H_2_O_2_ are shown. (B) Fluorescence micrographs showing cells of WT (PY79), Δ *ypsA* (RB42), and Δ *ypsA* complemented with a copy of inducible *ypsA* at an ectopic locus (RB160) grown with or without 1 mM H_2_O_2_ and stained with FM4-64 (membrane, red) and DAPI (DNA, blue). Arrow indicates aberrantly shaped cell. Scale bar: 1 μm.

### Increased production of YpsA inhibits cell division

Next, we examined *ypsA* overexpression phenotype. For this purpose, we constructed an otherwise WT-strain to ectopically express either *ypsA* or *ypsA-gfp* upon addition of inducer. Quantification of GFP fluorescence revealed that there was 3-fold overproduction of YpsA-GFP in the presence of inducer (2415 ± 1296 arbitrary units; n =50) when compared to YpsA-GFP produced under the control of its native promoter (732 ± 692 arbitrary units; n =50). We then monitored the cell morphology of cells overproducing YpsA or YpsA-GFP. To our surprise, as shown in Fig. 3, when compared to the cell lengths of inducible strains grown in the absence of inducer [YpsA: 2.92 ± 0.81 μm (Fig. 3A); YpsA-GFP: 3.89 ± 0.98 μm (Fig. 3C); n =100], cells overproducing YpsA or YpsA-GFP appeared elongated [YpsA: 8.92 ± 4.89 μm (Fig. 3B); YpsA-GFP:9.57 ± 4.99 μm (Fig. 3D); n =100] implying cell division is inhibited by YpsA. Also, this result indicated that the fluorescent protein tagged fusion of YpsA is functional. Tracking fluorescence of YpsA-GFP showed that YpsA assembles into discrete foci (Fig. 3D). Time-lapse microscopy conducted at 2 min interval for 10 min revealed that YpsA foci are highly dynamic (Fig. 3I-L). Since YpsA-GFP retains fluorescence as a focus and that the foci are mobile, and focus disruption occurs in some YpsA mutants (Fig. 6), we conclude that the foci are not artifacts of non-functional misfolded aggregates.

**Figure 3.**
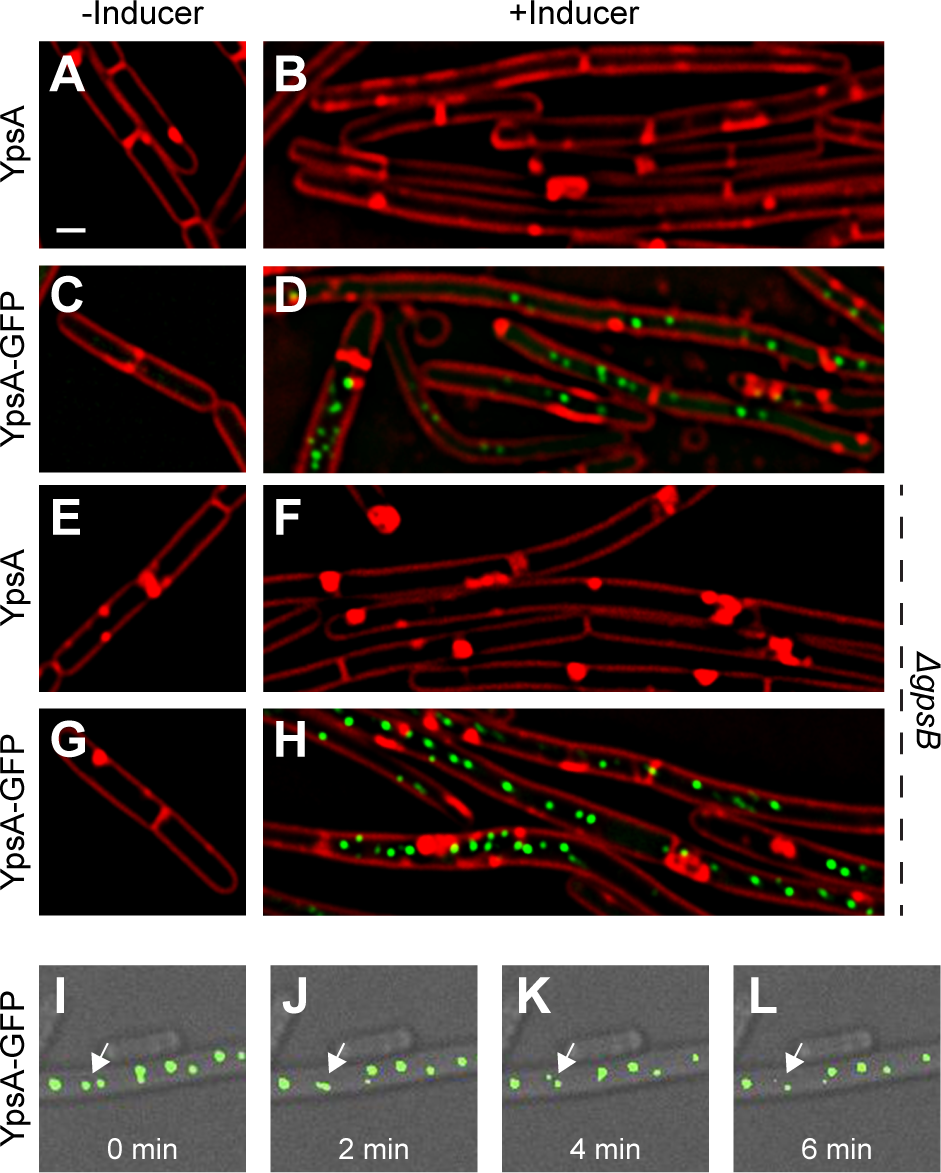
Elevated production of YpsA or YpsA‐GFP leads to inhibition of cell division. (A-D) Morphology of cells containing inducible *ypsA* (GG82) or *ypsA-gfp* (GG83) grown in the absence of inducer IPTG (A and C) or in the presence of inducer (B and D). (E-H) Cells morphology of strains lacking *gpsB* and containing either inducible *ypsA* (RB43) or *ypsA-gfp* (RB44) grown in the absence (E and G) or presence (F and H) of inducer. (I-L) Timelapse micrographs of *ypsA-gfp* expressing cells (GG83) and time intervals are indicated at the bottom. Arrow indicates foci that are mobile. DIC (gray) and fluorescence of membrane dye (FM4-64; red), GFP (green) are shown. Scale bar: 1 μm.

Since genes coding for YpsA and GpsB, a known cell division protein, are in a syntenous relationship we aimed to test whether YpsA overproduction-mediated filamentation is dependent on GpsB. As shown in Figs. 3E-H, cells lacking *gpsB* also formed filaments upon overexpression of *ypsA* or *ypsA-gfp*, suggesting that YpsA-mediated cell division inhibition is independent of GpsB.

Typically, filamentation is a result of impaired FtsZ ring assembly. To test whether FtsZ ring assembly is affected by YpsA overproduction, we engineered a strain that constitutively produces FtsZ‐GFP (28, 29), to also produce either *ypsA* or *ypsA-mCherry* under the control of an inducible promoter. In FtsZ-GFP producing otherwise WT cells, the cell length appeared normal and FtsZ assembled into FtsZ rings at mid-cell in 90% of the cells (Fig. 4A and 4B; top panels). In the strain capable of producing both FtsZ-GFP and YpsA or YpsA-mCherry, when cells were grown in the absence of inducer, FtsZ-GFP localization appeared similar to the control strain (Fig. 4A and 4B; middle panels). In striking contrast, when cells were grown in the presence of inducer FtsZ-GFP did not assemble into rings and instead appeared diffused (Fig. 4A and 4B; bottom panels).

**Figure 4.**
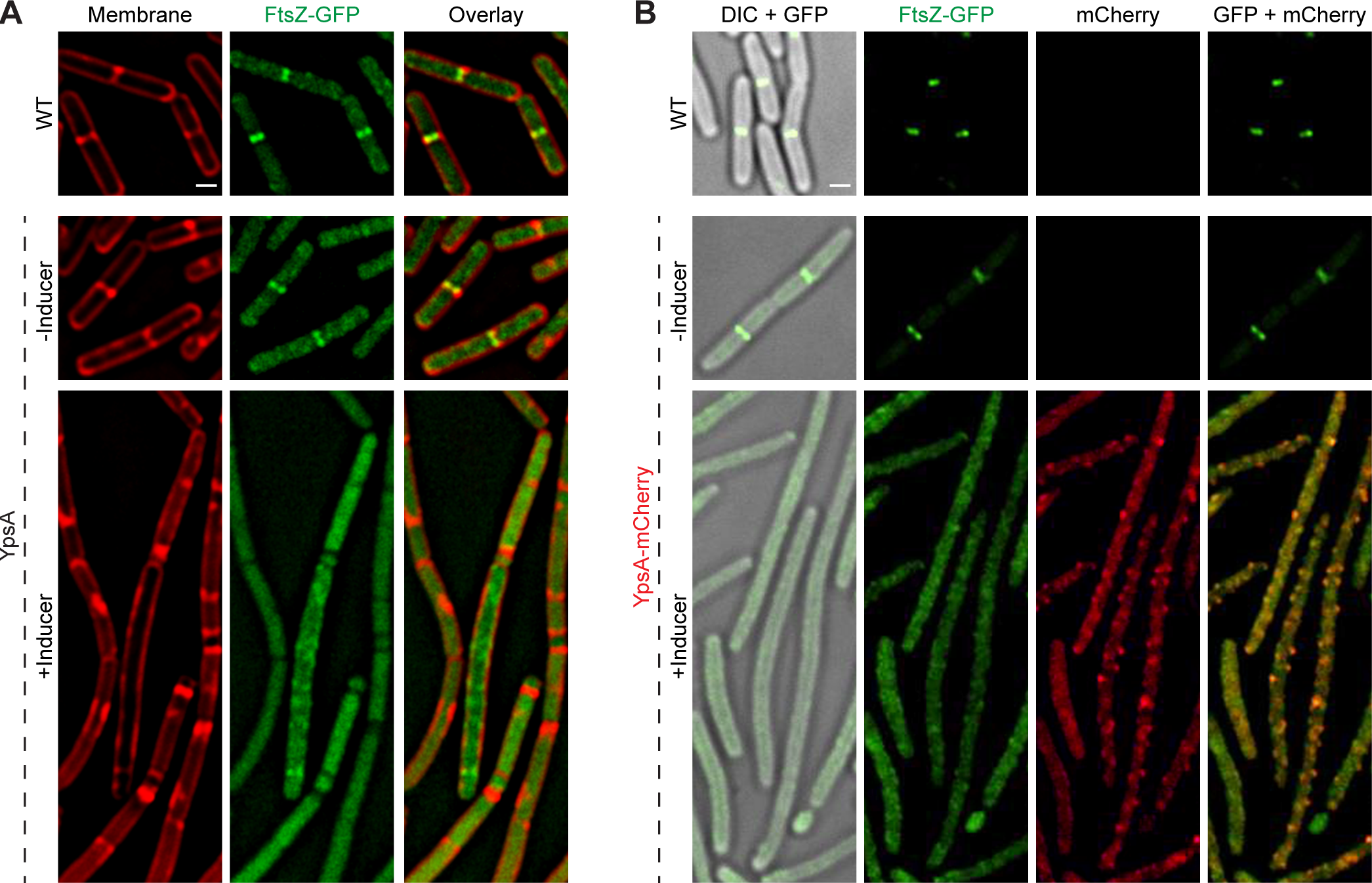
YpsA inhibits FtsZ ring assembly. (A) Fluorescence micrographs of cells that either constitutively produce FtsZ-GFP in otherwise wild type strain (PE92; top panel) and cells that constitutively produce FtsZ-GFP and additionally harbor a copy of inducible *ypsA* (RB15) grown in the absence (middle panel) or presence of inducer IPTG are shown. Fluorescence of FM4-64 membrane dye (red) and GFP (green) are shown.(B) Cellular morphologies of cells that constitutively produce FtsZ-GFP (PE92) and cells that additionally contain a copy of inducible *ypsA-mCherry* (RB97) grown in the absence (middle panel) or presence of inducer are shown. DIC (gray) and fluorescence of GFP (green) and mCherry (red) are shown. Scale bars: 1 μm.

### Filamentation is dependent on glucose availability

As *cotD*, which codes for a spore coat protein is immediately upstream of *ypsA* (Fig. S1A), we were curious to see if *ypsA* has any role in sporulation. To address this, we performed a sporulation assay using Casein Hydrolysate (CH)-based growth medium and Sterlini-Mandelstam sporulation medium in triplicates (30). The average sporulation frequency of Δ*ypsA* strain was 176% relative to WT (100%), which is a modest 2-fold increase in frequency suggesting YpsA has no appreciable role in sporulation. To study whether YpsA overproduction-mediated filamentation impairs sporulation, we conducted a similar sporulation assay and found that cells overexpressing *ypsA* (98%) or *ypsA-gfp* (127%) also displayed sporulation frequency similar to WT. To fully comprehend how filamentous cells achieve WT-like sporulation efficiency, we observed the cell morphology of *ypsA* overexpressing cells grown in the presence of inducer in CH medium using fluorescence microscopy. The cell lengths of *ypsA* or *ypsA-gfp* overexpressing cells appeared similar when grown with or without the inducer [YpsA: 2.85 ± 0.72 μm (Fig. 5A) vs 3.23 ± 0.93 μm (Fig. 5B); YpsA-GFP: 3.01 ± 0.59 μm (Fig. 5E) vs 3.51 ± 1.21 μm (Fig. 5F); n =100], unlike what we observed when cells were grown in LB medium (compare Figs. 5AB and Figs. 5EF with Figs. 3A-D). Although cells were not filamentous, YpsA foci still formed in CH (Fig. 5F).

**Figure 5.**
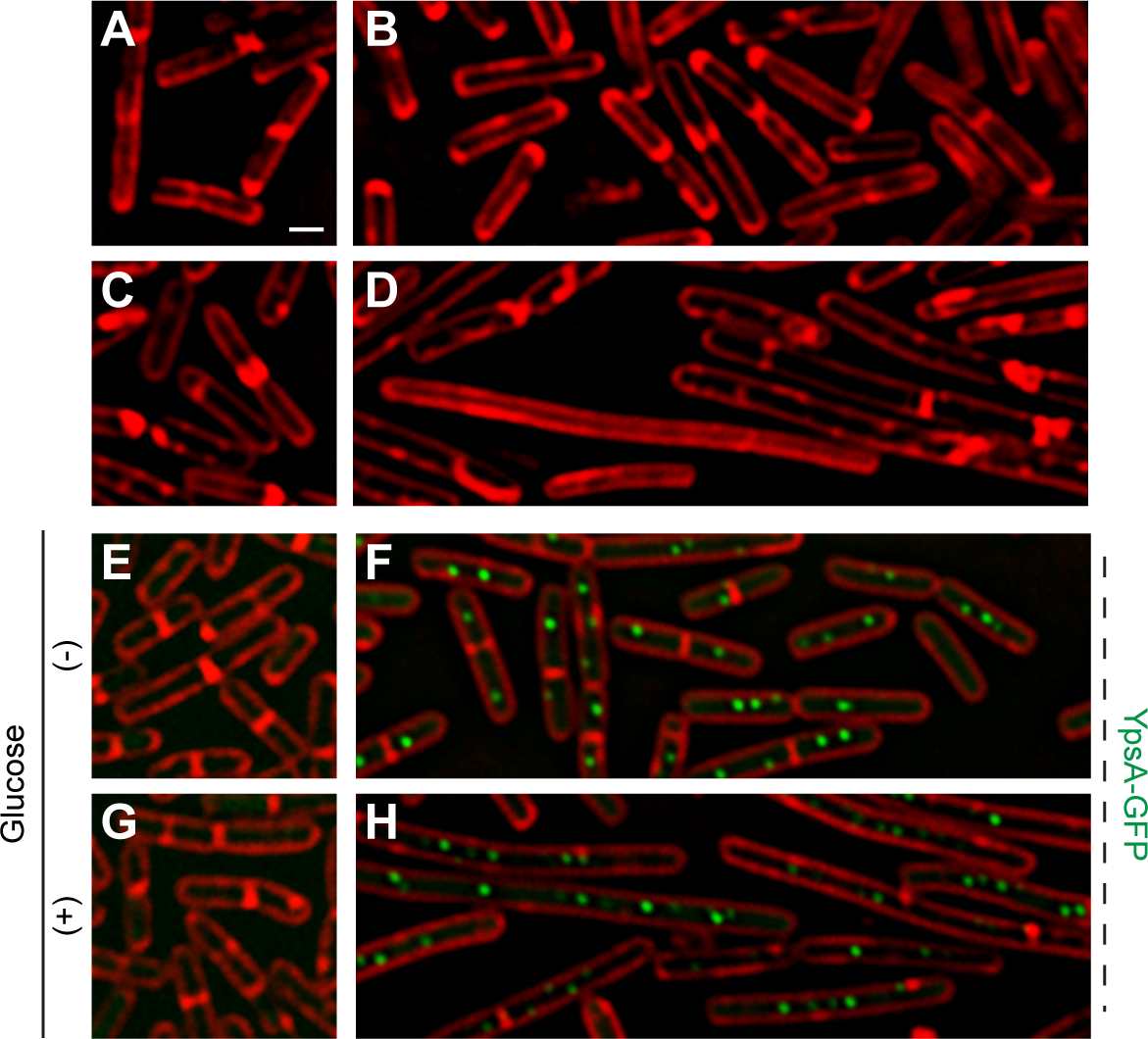
YpsA-mediated cell division inhibition is dependent on glucose availability. (AD) Fluorescence micrographs of cells containing inducible *ypsA* (GG82) or *ypsA-gfp* (GG83) grown in the absence of (A and C; E and G) or in the presence of inducer IPTG (B and D; F and H). The cells were grown either in the absence (A and B; E and F) or presence (C and D; G and H) of 1% D-glucose. Fluorescence of membrane dye (FM4-64; red), GFP (green) are shown. Scale bar: 1 μm.

We hypothesized that lack of nutrients in CH compared to LB might be the reason for lack of filamentation. To test our hypothesis, we externally added 1% glucose to the CH medium. Intriguingly, cells grown in CH in the presence of glucose and inducer to overproduce YpsA or YpsA-GFP lead to filamentation [YpsA: 3.31 ± 0.79 μm (Fig. 5C) vs 6.49 ± 2.95 μm (Fig. 5D); YpsA-GFP: 3.34 ± 0.94 μm (Fig. 5G) vs 9.48 ± 4.05 μm (Fig. 5H); n =100], suggesting that filamentation is dependent on metabolic status: specifically glucose availability in this case.

### Identification of amino acid residues important for YpsA function

Aided by the crystal structure and computational analysis of the YpsA family of SLOG domains we identified the conserved residues that are predicted to be important for maintaining the function of YpsA (Fig. 1B; see arrows). We performed site-directed mutagenesis of two of these key residues and generated GFP-tagged *ypsA* variants G53A and E55Q. We also generated other mutants to more generally explore YpsA function namely, G42A, E44Q, W45A, W57A, or W87A. We ensured that all mutants were stably produced through immunoblotting (Fig. 6B). Microscopic examination revealed that all YpsA variants except W57A were unable to trigger filamentation upon overexpression (Fig. 6A), suggesting that YpsA function is compromised in all these cases. We also noticed that G53A, E55Q, W45A, and W87A mutants displayed impaired ability to form foci. This is consistent with the observation that the first two of these mutations disrupt the conserved, predicted nucleotide-binding site of the YpsA family (21), and the latter two likely disrupt a key strand and helix of the Rossmannoid fold.

**Figure 6.**
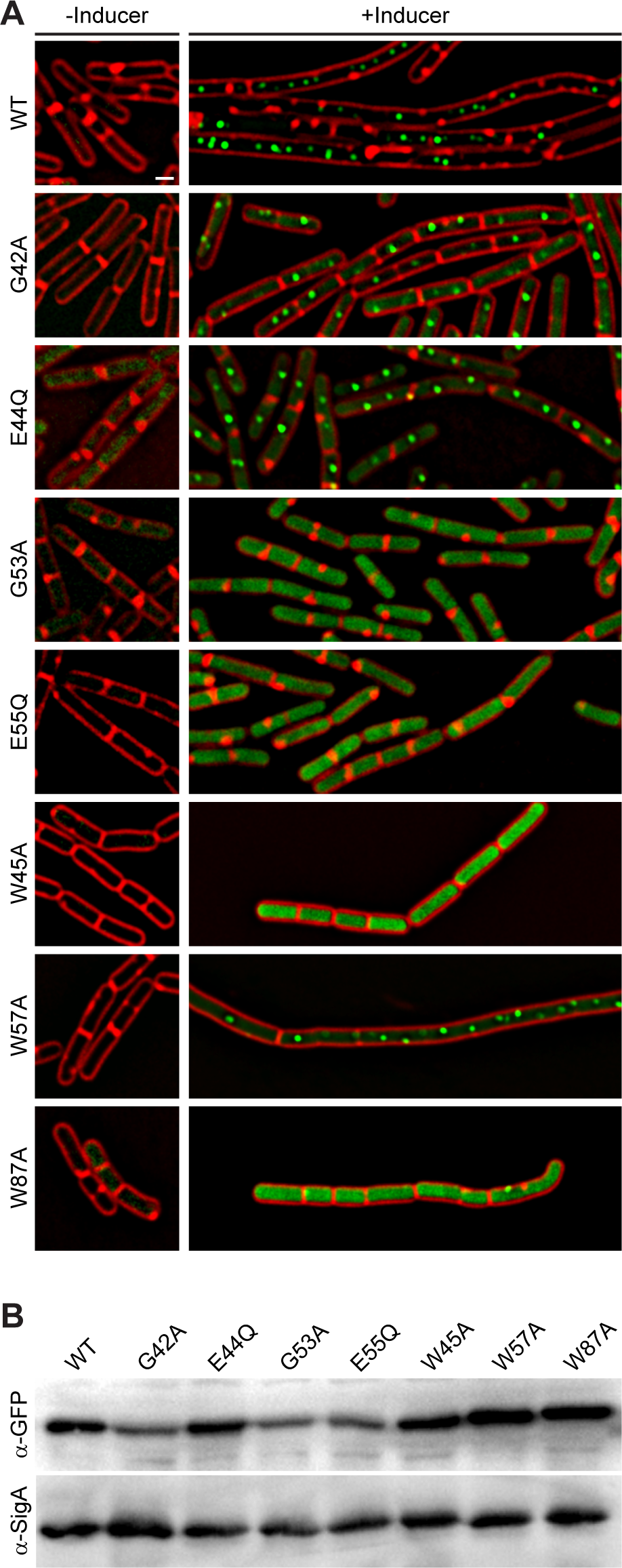
Site-directed mutagenesis reveals key residues in YpsA. (A) Cell morphologies of YpsA-GFP (WT; GG83) and GFP-fusions of G42A (RB119), E44Q (RB115), G53A (RB120), E55Q (RB116), W45A (RB35), W57A (RB26), and W87A (RB37) are shown. The cells were grown either in the absence (left panels) or presence (right panels) of inducer IPTG. Fluorescence of membrane stain FM4-64 (red) and GFP (green) are shown. Scale bar: 1 μm. (B) Production of GFP-tagged YpsA variants were detected by immunoblot of cell extracts of strains shown in (A) grown in the presence of inducer using anti-GFP and corresponding anti-SigA (loading control) antisera.

### Putative interaction partners of YpsA

To understand the role of YpsA via identifying its potential interaction partners, we conducted FLAG-immunoprecipitation using YpsA-FLAG and YpsA-GFP-FLAG constructs as baits. Untagged YpsA served as our negative control. After confirming the enrichment of proteins in the eluate fractions through silver staining and anti-Flag immunoblotting, the samples were submitted for protein identification via mass spectrometry. To identify proteins that specifically interact with YpsA, we eliminated all proteins that also appeared in our negative control, as they are likely non-specifically bound proteins and retained only proteins that were present specifically in both YpsA-FLAG and YpsA-GFP-FLAG eluates. A selective list of protein interaction partners of YpsA is shown in Table.1. The entire list is provided in Table. S2. In addition to our bait, FLAG-tagged versions of YpsA and presumably native copies of YpsA due to self-assembly, we noticed many proteins whose genes are under nutrient availability-sensing CcpA (31) or CodY (32) or AbrB (33) regulon(s) in our IP results. Interestingly, several of the proteins that associate with YpsA bind NAD or its derivatives and/or play a role in redox-sensing. However, given that these are abundant metabolic enzymes we cannot be sure of the significance of these interactions at this time.

**Table 1.**
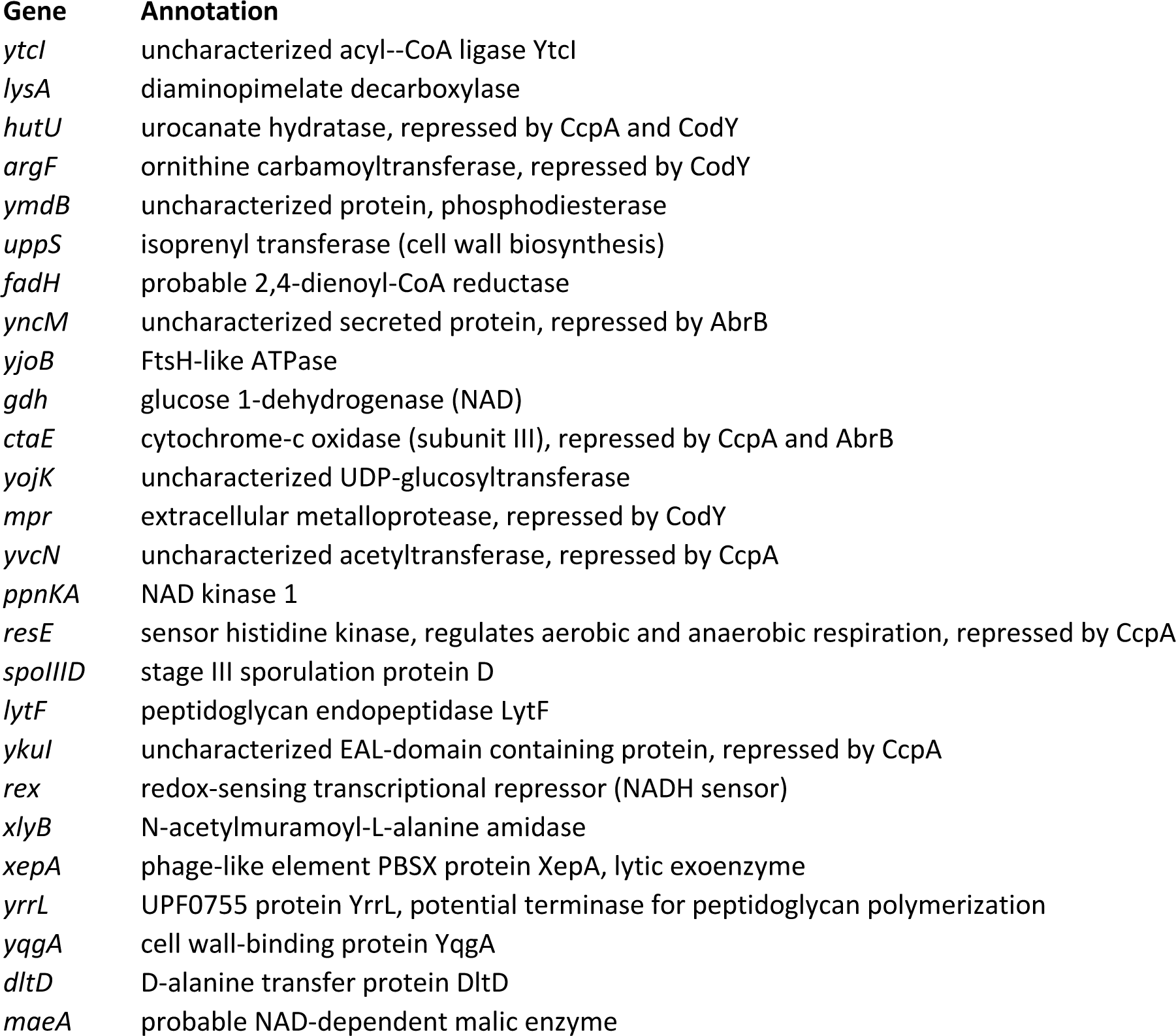
Selective list of putative interaction partners of YpsA

### Overproduction of YpsA inhibits cell division in *S. aureus*

To investigate if the role of YpsA is conserved in other Firmicutes, we chose to study the function of YpsA in *S. aureus*. Cells lacking intact *ypsA* in *S. aureus* (34), are viable and their cell morphology appear similar to WT control suggesting *ypsA* is not an essential gene (Fig. 7AB). Next, we placed *S. aureus ypsA* (*ypsA ^SA^*) under the control of xylose-inducible promoter in a *S. aureus* plasmid vector. As shown in Fig. 6, the cell diameter of WT control (0.86 ± 0.18 μm; n =100; Fig. 7A) and vector control strain grown in the absence of inducer (0.99 ± 0.21 μm; n =100; Fig. 7C) and presence of inducer (1.12 ± 0.21 μm; n =100; Fig. 7D), resembled inducible *ypsA ^SA^* strain grown in the absence of inducer (1.19 ± 0.30 μm; n =100; Fig. 7E). Interestingly, cells overexpressing *ypsA^SA^* were unable to undergo septation and displayed clear cell enlargement (1.72 ± 0.37 μm; n =100; Fig. 7F), a telltale sign of cell division inhibition in this organism. Thus, the function of YpsA in inhibiting cell division is conserved in *S. aureus,* and possibly in other Firmicutes which code for it despite the differences in their cell-morphology.

**Figure 7.**
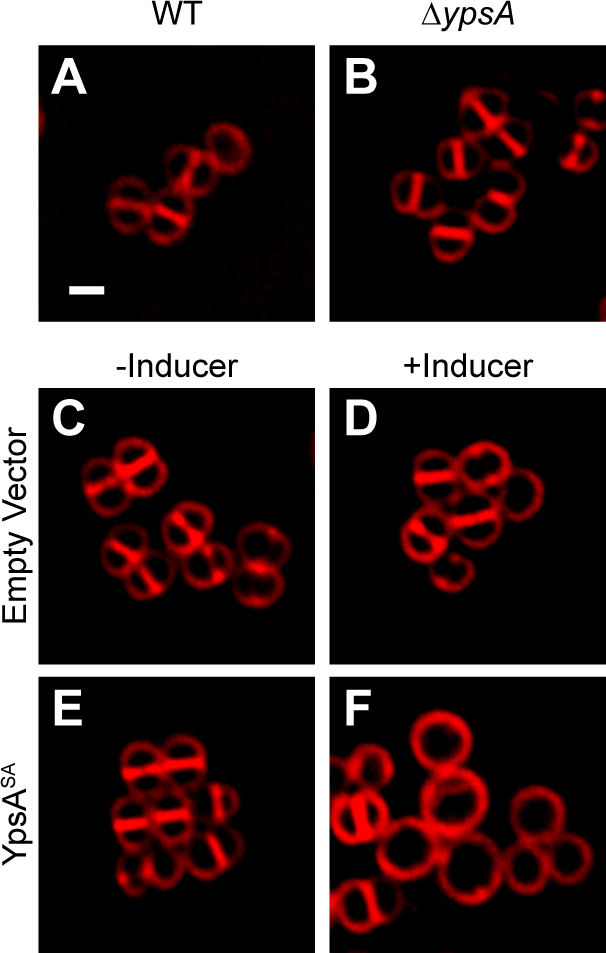
Production of YpsA^SA^ inhibits cell division in *S. aureus*. (A-B) Fluorescence micrographs of wild type (SH1000; A), transposon-disrupted *ypsA* (RB162; B) strains. (C-F) Morphologies of SH1000 cells harboring plasmid encoded xylose-inducible copy of *ypsA ^SA^* (pRB36; E and F) or empty vector (pEPSA5; C and D) grown in the absence (C and E) and presence (D and F) of inducer are shown. Membranes were visualized using FM4-64 dye (red). Scale bar: 1 μm.

## DISCUSSION

Bacterial cell division is a highly regulated process and many division factors have already been characterized especially in model organisms *E. coli* and *B. subtilis*. Yet, cell division is only mildly affected even in the absence of a combination of known division regulators in these organisms (12), thus predicting the presence of other proteins that could affect the cell division process. Here, we discuss the role of YpsA, a protein of hitherto unknown function conserved in diverse Firmicutes. We show that YpsA offers protection against oxidative stress. However, the precise mechanism of how this is achieved remains to be elucidated. Next, we show that YpsA overproduction leads to impaired FtsZ ring assembly and ultimately cell division inhibition.

It has been reported that *cotD-ypsA* transcriptional unit is repressed by the regulator essential for entry into sporulation, Spo0A (35), which binds to a region upstream of *cotD* (36). It has been shown that *cotD* is also repressed by a late stage sporulation-specific transcriptional regulator, SpoIIID (37). Both *cotD* and *ypsA* transcripts are at similar levels in various growth conditions except in those that promote sporulation [Fig. S1B; (26, 27)]. The function of CotD during normal growth, if any, needs to be evaluated. It has been reported that *cotD* level increases in a concentration-dependent manner in response to antibiotic treatment (38). In this report we show that cells lacking *ypsA* or overexpressing *ypsA* show no obvious sporulation defect and that YpsA-mediated cell division inhibition is dependent on glucose availability. Other reports exist that show a clear connection between glucose availability and cell division (1, 39). One such factors that inhibit cell division depending on the presence of glucose is UgtP which is a UDPglucose diacylglycerol glucosyltransferase (40). However, as shown in Fig. S2, cell lacking *ugtP* also undergo filamentation upon increased production of YpsA, suggesting that cell division inhibition by YpsA is independent of UgtP.

YpsA mutants generated based on the crystal structure and sequence analysis revealed the importance of certain key residues for YpsA function (Fig. 6). Interestingly, G53 and E55 of *B. subtilis* YpsA which form a conserved signature GxD/E motif, are predicted to be important for substrate-binding in YpsA clade of proteins in SLOG superfamily [Figure 1B; (21)]. Since foci-formation was disrupted in both G53A and E55Q mutants, it is plausible substrate-binding allows for multimeric complex formation. It is noteworthy that mutants such as G42A and E44Q which are able to form foci, therefore likely bind substrate, lack the ability to elicit filamentation. Also, YpsA-GFP overproducing cells grown in CH medium were able to form foci but unable to induce filamentation (Fig. 5F). These observations support a model in which substrate binding by YpsA is a prerequisite for cell division inhibition but substrate binding alone is not sufficient to induce filamentation, assuming foci formation is indicative of substrate binding. It is possible YpsA executes cell division inhibition function through interactions with other protein partners.

Consistent with this our pull-down assay identified multiple putative interaction partners of YpsA, including multiple NAD-binding proteins. Interestingly, the connection between NAD or its derivative ADP-ribose and the members of SLOG superfamily of proteins that belong to YpsA clade has been previously suggested (21). Given that ADP-ribosylation affects FtsZ polymerization (41, 42), and YpsA is in close association with biological conflict systems and phosphoribosyl transferases (Fig. 1B), it is possible that YpsA-mediated inhibition of cell division may involve ADP-ribosylation. Similarly, oxidative stress protection provided by YpsA might involve sensing or binding NAD or its derivatives as well. The link between metabolism of nicotinamide nucleotide, glucose availability, and oxidative stress has been reported previously (43, 44).

Lastly, we show that YpsA in another Firmicute, *S. aureus*, also inhibits cell division, hinting at a conserved role for YpsA in these Gram-positive organisms. In *B. subtilis*, a prophage associated protein of unknown function, YoqJ, also belongs to the YpsA family (Fig. 1B). Given that there are clear examples of phage proteins affecting bacterial cell division (45-48), it would be interesting to see if YoqJ also influences cell division. Although the GxD/E motif is conserved in YoqJ, several residues we identified to be essential in YpsA are not conserved in YoqJ (Fig. 1B and Fig. 6). The Firmicutes-specific conserved gene coupling between *ypsA* and *gpsB* starkly contrasts the diversity of the gene neighborhoods found in other branches of YpsA family phylogenetic tree. Superposition of conserved gene-neighborhoods onto the phylogenetic tree (Fig. 1A) revealed a stark compartmentalization in conserved genome contexts. The *ypsA* and *gpsB* gene coupling is found only in one of the four major branches in the tree. Each of three others display distinct conserved neighborhood proclivities: 1) a branch where YpsA couples strongly in a gene pair relationship with a phosphoribosyltransferase (PRTase) domain, 2) a branch where YpsA is found in scattered associations with various components of NAD processing and salvage pathways, and 3) a diverse collection of contexts across a broad class of bacterial lineages representative of the aforementioned nucleotide-centered biological conflict systems, where YpsA is likely to act in nucleotide signal-generation or nucleotide-sensing [Fig. 1B; (21)]. These observations suggest that the *B. subtilis* YpsA may have acquired a more institutionalized role in cell division within the Firmicutes phylum. Nevertheless, understanding the precise biochemical mechanism by which *B. subtilis* YpsA executes its function would potentially shed light on the more general function of YpsA across a wide range of organisms and biological conflict systems.

## MATERIALS AND METHODS

### Strain construction and general methods

All *B. subtilis* strains used in this study are isogenic derivatives of PY79 (49). See table S1A for strain information. Overproduction of YpsA was achieved by PCR amplifying *ypsA* using primer pairs oP106/oP108 (see table S1B for oligonucleotide information) and ligating the fragment generated cut with SalI and NheI with IPTG-inducible *amyE* locus integration vector pDR111 (D. Rudner) also cut with SalI and NheI and the resulting plasmid was named pGG27. To construct a GFP fusion, *ypsA* fragment that was amplified with primer pairs oP106/oP107 and digested with SalI and NheI was ligated with *gfp* fragment generated with oP46/oP24 and cut with NheI/SphI and cloned into pDR111 digested with SalI/SphI resulting in plasmid pGG28. The G42A, E44Q, W45A, G53A, E55Q, W57A, and W87A mutations were introduced using the QuikChange site-directed mutagenesis kit (Agilent) using pGG28 as template. *ypsA-3xflag* was constructed via two step PCR using pGG27 as a template. Round one PCR was completed using primers oP106 and oP291. The PCR product from round one was then used as a template for round two PCR, which was completed using primer pairs oP106 and oP292. The final PCR product was then cloned into pDR111 using SalI and NheI restriction sites, making plasmid pRB33. Similarly, *ypsA-gfp-3xflag* was constructed via two step PCR using pGG28 as a template. Round one PCR was completed using primers oP106 and oP349. The PCR product from round one was then used as a template for round two PCR, which was completed using primers oP106 and oP350. The final PCR product was then cloned into pDR111 using SalI and SphI restriction sites, making plasmid pRB34. The engineered plasmids were then used to introduce genes of interest via double crossover homologous recombination into the *amyE* locus of the *B. subtilis* chromosome. Expression of *ypsA-his* in BL21-DE3 *Escherichia coli* cells was achieved by PCR amplifying *ypsA-his* with primer pairs oRB9 and oRB33, and cloning into XbaI and BamHI resticition sites of pET28a, producing plasmid pRB21. YpsA-his was purified using standard protocol involving nickel column-based affinity chromatography. To produce *S. aureus* YpsA in *S. aureus* strain SH1000, *ypsA ^SA^* fragment (PCR amplified with oRB27/oP314 primer pairs) was cloned into xylose-inducible pEPSA5 plasmid using EcoRI and BamHI restriction sites (50), generating plasmid pRB36. Plasmids were first introduced into *S. aureus* RN4220 via electroporation, and then transduced into SH1000 (18).

### Media and culture conditions

Overnight *B. subtilis* cultures grown at 22 °C in Luria-Bertani (LB) growth medium were diluted 1:10 into fresh LB medium and grown to mid-logarithmic growth phase (OD_600_ = 0.5), unless otherwise stated. Expression of genes under IPTG-controlled promoter was induced by addition of 1 mM IPTG (final concentration) to the culture medium unless noted otherwise. Overnight *S. aureus* cultures were grown at 22°C in tryptic soy broth (TSB) supplemented with 15 µg/ml chloramphenicol and/or 5 µg/ml erythromycin where required for plasmid maintenance. Cultures were then diluted 1:10 into fresh medium containing appropriate antibiotics and grown to mid-logarithmic growth phase (OD_600_ =0.5), unless otherwise stated. Expression of genes under xylose-controlled promoter was induced by the addition of 1% xylose when required.

### Sporulation assay

Sporulation assay was conducted using resuspension protocol as described previously (30). Briefly, overnight cultures of *B. subtilis* cells were grown in LB medium at 22°C, were diluted 1:10 in fresh casein hydrolysate medium (CH, KD Medical) and grown to mid-log phase twice before culture was resuspended in Sterlini-Mandelstam sporulation medium (SM, KD Medical) to induce sporulation (51). Growth in CH medium and entry into sporulation in SM medium were monitored via fluorescence microscopy. Total viable cell counts (CFU/ml prior to heat treatment) and spore counts (CFU/ml after incubation at 80°C for 10 min) were obtained for calculating sporulation frequency (spore count/viable count).

### Disc diffusion assay

All disc diffusion assays were completed on LB agar plates. Strains PY79 and RB42 were grown until OD_600_=0.5, and then 100μl of each culture was added to the respective plates. Briefly, 15μl of 1M hydrogen peroxide was added to 7mm Whatman paper discs, which were then placed equidistant from each other on top of the inoculated media. A disc containing no hydrogen peroxide was used as a negative control. Plates were then incubated overnight at 37°C. The diameter of the disc (7mm) was subtracted for the zone of inhibition measurements.

### Immunoprecipitation and mass spectrometry

The YpsA-FLAG immunoprecipitation was performed using FLAGIPT1 immunoprecipitation kit (Sigma-Aldrich) as described previously (52). Briefly, 1 ml cell lysates of cells harvested from 20 ml LB culture induced with 1 mM IPTG (final concentration) grown for 2 h post-induction to produce FLAG-tagged proteins or untagged negative control were generated by sonication. Cell extracts were then incubated overnight with 50 µl anti-FLAG M2 affinity beads supplied by the manufacturer. The beads were then washed 3 times with 1x wash buffer and the supernatant was removed by pipetting. Proteins bound to the beads were stripped by adding 80 μl of 2x sample buffer supplied by the manufacturer and heating at 100 °C for five minutes. The supernatants were collected and subjected to SDS-PAGE analysis prior to mass spectrometry. Western blot using anti-Flag antibody (Invitrogen) was used to detect Flag-tagged proteins in all fractions collected.

For mass spectrometry, protein extracts were separated by SDS-PAGE and silver-stained for visualization. The gel was divided into 3 fractions, and each gel section was minced and de-stained before being reduced with dithiothreitol (DTT), alkylated with iodoacetamide (IAA), and finally digested with Trypsin/Lys-C overnight at 37 °C. Peptides were extracted using 50/50 acetonitrile (ACN)/H_2_O/0.1% formic acid and dried in a vacuum concentrator. Peptides were resuspended in 98%H_2_O/2%ACN/0.1% formic acid for LC-MS/MS analysis. Peptides were separated using a 50 cm C18 reversed-phase HPLC column (Thermo Scientific) on an Ultimate3000 UHPLC (Thermo Scientific) with a 60 min gradient (2-32% acetonitrile with 0.1% formic acid) and analyzed on a hybrid quadrupole-Orbitrap mass spectrometer (Q Exactive Plus, Thermo Fisher Scientific) using data-dependent acquisition in which the top 10 most abundant ions are selected for MS/MS analysis. Raw data files were processed in MaxQuant [19029910] and searched against the current UniprotKB *Bacillus subtilis* 168 protein sequence database. Search parameters include constant modification of cysteine by carbamidomethylation and the variable modification, methionine oxidation. Proteins are identified using the filtering criteria of 1% protein and peptide false discovery rate.

### Microscopy

Aliqouts containing 1 ml of culture (*B. subtilis* and *S. aureus*) were washed in phosphate buffered saline (PBS) and then resuspended in 100 μl of PBS containing 1 μg/ml FM4-64 (membrane stain) and/or 2 μg/ml DAPI (DNA stain). For imaging, 5 μl of sample was then spotted onto a glass bottom dish (MatTek) and it was covered with an 1% agarose pad made with sterile water. Still imaging was completed at room temperature. For time-lapse microscopy, 5 μl aliquots of culture were spotted onto a glass bottom dish, and the sample was covered with 1% agarose pad made with LB culture medium. Agarose pads were supplemented with FM4-64 and/or DAPI to stain the cell membrane and DNA respectively during the course of data collection, and inducer where required to induce the expression of desired genes. Microscopy was performed using GE Applied Precision DeltaVision Elite deconvolution fluorescence microscope equipped with a Photometrics CoolSnap HQ2 camera and environmental chamber. Typically, 17 planes (Z-stacks) every 200 nm was acquired of all static image data sets and 5 planes every 200 nm was acquired for time-lapse microscopy to minimize phototoxicity. The images were then deconvolved using SoftWorx software provided by the manufacturer.

### Sequence Analysis

YpsA sequence similaritiy searches were performed using the PSI-BLAST program (53) against the non-redundant (NR) database of the National Center for Biotechnology Information (NCBI). Multiple sequence alignments were built by the MUSCLE and KALIGN programs (54, 55), followed by manual adjustments on the basis of profile– profile and structural alignments. Genes residing in conserved neighborhoods were identified through clustering carried out with the BLASTCLUST program (ftp://ftp.ncbi.nih.gov/blast/documents/blastclust.html). Phylogenetic analysis was conducted using an approximately-maximum-likelihood method implemented in the FastTree 2.1 program under default parameters (56), and resulting trees were visualized initially in the FigTree program [http://tree.bio.ed.ac.uk/software/figtree/].

## ACKNOWLEDGEMENTS

We thank our lab members and K. Ramamurthi for comments on the manuscript; University of South Florida (USF) Department of Cell Biology, Microbiology and Molecular Biology - Mass Spectrometry Core Facility for proteomics support. This work was funded by a start-up grant from USF (P.J.E.) and the National Institutes of Health (NIH) grant (R01GM128037; P.J.E.). A.M.B and L.A. are supported by the Intramural Research Program of the National Library of Medicine, NIH, USA.

## AUTHOR CONTRIBUTIONS

R.S.B. and P.J.E. designed the study. R.S.B, G.G., S.S.A., M.H., M.W., and P.J.E. constructed strains and performed experiments. A.M.B. and L.A. conducted bioinformatics analysis. R.S.B., M.W., A.M.B, L.A., and P.J.E. analyzed data. R.S.B. and P.J.E. wrote the paper. All authors read and commented on the final manuscript.

## Supplemental data

**Figure S1.** (A) Cartoon representation of *ypsA* gene neighborhood. (B) Transcript levels of *cotD* and *ypsA* in *B. subtilis* at various growth conditions (26, 27).

**Figure S2.** Cell morphologies of inducible *ypsA* cells (GG82) grown in the absence (A) or presence (B) of inducer. Also shown are the cell morphologies of inducible *ypsA* in a strain lacking *ugtP* (RB212) grown in the absence (C) or presence (D) of inducer.

**Table S1.** Strains and oligonucleotides used in this study

**Table S2.** Putative interaction partners of YpsA

